# Narcissistic and Antisocial Personality Traits are both encoded in the Triple Network: Connectomics evidence

**DOI:** 10.1101/2024.12.03.626524

**Authors:** Khanitin Jornkokgoud, Richard Bakiaj, Peera Wongupparaj, Remo Job, Alessandro Grecucci

## Abstract

The neural bases of narcissistic and antisocial traits are still under debate. One intriguing question is whether these traits are encoded within the so-called triple network e.g. the default mode (DMN), salience (SN), and fronto-parietal (FPN) networks, and whether these traits affect the same networks in a similar manner. Connectome-based analyses were conducted on resting-state scans from 183 participants, examining regional and global graph-theoretic metrics in the DMN, SN, and FPN, with the visual and sensorimotor networks as controls. Our findings revealed a clear involvement of the triple network in narcissistic and antisocial traits, confirming a shared neural substrate for the two traits. Both traits were negatively predicted by the anterior cingulate cortex of the SN, possibly indicating less awareness of dangers and more proneness to engage in risky behaviors. Additionally, both traits were positively predicted by the lateral prefrontal cortex of the FPN, suggesting augmented strategic thinking to manipulate others and increased planning skills to achieve personal goals. Besides similarities, there were also some differences. Specific hubs of the DMN were positively associated with narcissism but negatively related with antisocials, possibly explaining their differences in self-reflection and thinking about the self, largely present in the former, but usually reduced in the latter. These results extend previous findings on the involvement of the triple network in personality disorders and suggest both common and different mechanisms underlying narcissistic and antisocial traits. As such, these findings could pave the way for developing potential biomarkers of personality pathology and identify neurostimulation intervention targets.

## INTRODUCTION

Narcissism exists on a continuum, ranging from subtle, sub-clinical traits to the pathological condition of Narcissistic Personality Disorder (NPD), with each manifestation impacting mental health and interpersonal relationships to varying degrees (Nenadić et al., 2021). This continuum reflects alterations in psychological components such as self-esteem regulation, self-coherence, and empathy (Miller et al., 2011; Ronningstam, 2011). At one extreme of the continuum lies the NPD, which is characterized by overt expressions of dominance, inflated self-importance, and anger. Individuals with this grandiose version of narcissism often exhibit behaviors marked by ego inflation and a dominant interpersonal style (Miller et al., 2011). A more subtle version of NPD exists e.g. vulnerable narcissism,vulnerable narcissism is associated with insecurity, hypersensitivity, and feelings of shame tend to mask their underlying insecurities through defensive negativity (Miller et al., 2011). Apart from fully diagnosed disorder, it is important to emphasize that narcissistic traits can be present in the general population without necessarily meeting the criteria for a full disorder.

This observation is central to the ongoing debate about how personality disorders should be diagnosed. Currently, two primary systems are used for diagnosing mental disorders: the fifth edition, text revision, of the Diagnostic and Statistical Manual of Mental Disorders (DSM-5-TR) and the eleventh revision of the International Classification of Diseases (ICD-11). Both systems traditionally employ a categorical approach to differentiate personality disorders. However, these diagnostic categories often exhibit significant symptom overlap, potentially undermining their effectiveness and diagnostic accuracy (Day et al., 2024; George & Short, 2018; Hörz-Sagstetter et al., 2018). In response, a dimensional model has been proposed, suggesting that personality traits exists along a continuum from mild to severe (Ashton, 2013; Kotov et al., 2022). This dimensional model is incorporated in Section III of the DSM-5, which offers an alternative framework for diagnosis while still retaining the traditional categorical system (American Psychiatric Association, 2013). Conversely, the ICD-11 fully embraces a dimensional approach, proposing that personality disorders should be defined along a spectrum based on trait-specific features (Tyrer et al., 2015).

This shift towards a dimensional classification is also gaining traction in the field of neuroscience (Jornkokgoud et al., 2024; Jornkokgoud et al., 2023; Langerbeck et al., 2023). Indeed, neuroimaging research is shedding light on the neural underpinnings of personality traits (Grecucci et al., 2022; Jauk & Kanske, 2021), suggesting that identifying neurobiological markers may improve clinical practices by facilitating tailored treatments and predicting patient outcomes (Luo et al., 2022), potentially intervening before a trait becomes severe enough to warrant a categorical diagnosis. In this vein, current treatment modalities for personality disorders, emphasize the fact that they can be applied to individuals with sub-threshold diagnosis, or with other diagnostic categories where personality traits are play a role (Hawke & Provencher, 2011). Neuroscience applied to personality offered valuable insights into the underlying mechanisms of narcissism (DeYoung et al., 2022; Jauk et al., 2017). Structural MRI studies, for example, have identified significant variations in gray matter (GM) within the prefrontal cortex and insular regions, as well as in white matter (WM) microstructures in individuals with NPD, suggesting the presence of distinct neural signatures associated with this disorder (Jornkokgoud et al., 2024; Jornkokgoud et al., 2023; Nenadic et al., 2015). Furthermore, functional MRI investigations have revealed abnormal activation in several brain regions associated with narcissistic traits, offering valuable insights into the neurobiology of narcissism (Cao et al., 2022; Fan et al., 2011; Feng et al., 2018; Jauk et al., 2017). More recent studies have started clarifying the importance of functional connectivity, as opposed to structural connectivity, which yields more accurate behavioral predictions (Seguin et al., 2020). Resting-state functional connectivity (RSFC) analyses have identified key nodes, including the amygdala, prefrontal cortex (PFC), and the anterior cingulate cortex (ACC), as crucial areas involved in the neural network of narcissistic traits (Feng et al., 2018). Moreover, individuals with NPD exhibit altered RSFC patterns between brain regions in the default mode network (DMN) compared to healthy controls (Cao et al., 2022). This altered connectivity, particularly between cognitive networks and the DMN, has been shown to be a significant predictor of narcissistic traits (Cao et al., 2022; Jornkokgoud et al., 2024).

One intriguing, but poorly understood aspect is that narcissistic traits often combine with antisocial traits. Antisocial personality traits (APT) are, by definition, socially undesirable or antisocial, and in their most extreme form, strongly correlate with a clinical diagnosis of Antisocial Personality Disorder (ASPD) (American Psychiatric Association, 2013). The neuroimaging literature, including studies utilizing structural and functional MRI, as well as connectivity analyses, consistently supports the notion that psychopathy and antisocial traits are associated with impaired functioning across multiple brain regions (Blair, 2010; Gregory et al., 2012; Jiang et al., 2016; Jiang et al., 2013; Kumari et al., 2006; Kumari et al., 2013; Kumari et al., 2014; Tang et al., 2016). For instance, in structural MRI studies notable reductions in GM have been observed in several brain regions, including the frontopolar cortex, orbitofrontal cortex (OFC), frontal gyri, ACC, medial prefrontal cortex (MPFC), superior temporal gyrus, superior temporal sulcus, sensory-motor area, and rectal gyrus (Jiang et al., 2016; Johanson et al., 2020; Klaus et al., 2024; Kumari et al., 2014; Narayan et al., 2007; Raine et al., 2011).

In the same way, functional MRI studies during task-based paradigms have revealed decreased activity in key regions such as the left frontal gyrus, ACC, precuneus, thalamus, and dorsal ACC (dACC) during tasks including reward anticipation, Go/NoGo, Stroop, and n-back tasks (Barkataki et al., 2008; Kumari et al., 2006). Furthermore, resting-state fMRI in individuals with ASPD has demonstrated reduced activity in the posterior cerebellum and middle frontal gyrus, alongside increased activity in the inferior temporal gyrus, middle occipital gyrus, and inferior occipital gyrus (Kumari et al., 2006; Liu et al., 2014). Additionally, alterations in the topological organization of resting-state functional connectivity (RSFC) networks have been observed in antisocial individuals. These alterations include an increased clustering coefficient and decreased betweenness centrality in regions such as the medial superior frontal gyrus, precentral gyrus, Rolandic operculum, superior parietal gyrus, angular gyrus, and middle temporal pole (Tang et al., 2016). These findings highlight the complex neurobiological underpinnings of antisocial traits and their potential implications for understanding ASPD.

As anticipated, although separate disorders, narcissistic and antisocial personalities share commonalities. For example, they display some overlaps at the symptomatic level: dominance, inflated self-importance, anger outbursts, manipulation tendencies, and impulsivity. Also, it is common for both traits to be expressed in the same individuals. Previous research has indeed found that antisocial traits can predict narcissistic traits and vice versa (Jornkokgoud et al., 2023). Similarly, individuals with antisocial traits may be considered as exhibiting a more extreme form of narcissism (Kernberg, 1992; Miller et al., 2017). These observations have led some scholars to suggest new terminology to indicate this combination. For example, Paulhus and Williams have proposed the concept of the Dark triad of personality (Kernberg, 1992; Paulhus & Williams, 2002) that refers to individuals with high traits of Narcissism, Psychopathy, and Machiavellianism (e.g. the tendency to manipulate others) (Paulhus & Williams, 2002). Other researchers have proposed the concept of malignant narcissism, in which narcissistic, antisocial and paranoid traits combine to form the most extreme expression (Kernberg, 1989). No neural data are available yet on these assumed cobinations of traits.

Given the frequent co-occurrence of narcissistic and antisocial traits in certain individuals, which often leads to severe personality disturbances, it is crucial to investigate the common neural mechanisms underlying these traits. Previous research has highlighted the interaction of key brain networks, particularly the triple network, which includes the salience network (SN), central executive network or fronto-parietal network (FPN), and DMN in borderline personality (Doll et al., 2013; Krause-Utz et al., 2014; Langerbeck et al., 2023), but also in obsessive-compulsive personality disorder (OCPD) (Aguilar-Ortiz et al., 2020; Amiri et al., 2023; Coutinho et al., 2016; Doll et al., 2013; Krause-Utz et al., 2014; Tang et al., 2016). Nonetheless, previous studies focusing on macro networks in NPT or NPD have yielded limited results, as discussed in Jornkokgoud et al. (2024). Furthermore, collective evidencec suggests that alterations in the DMN, SN, and FPN (the triple network) may contribute not only to narcissistic traits but also to antisocial traits.

Of note the previous studies on the role of triple networks on personality traits have mainly used connectivity measures such as ROI-to-ROI, or seed-based connectivity, to quantify the communication between regions. However, the connectivity inside the triple network has remained largely unexplored. Brain network approaches allow us to establish a network that consists of elements and their pairwise interconnections, also referred to as nodes and edges (Faskowitz et al., 2022; Sporns, 2018). Specifically, the graph-based network method is a powerful tool for studying both functional and network connectivity, offering a detailed topological analysis of the brain’s network functionality (Farahani et al., 2019; Sporns, 2018). Moreover, this approach can provide novel insights into neurobiological mechanisms underlying cognition, behavior, and brain disorders (Farahani et al., 2019). The graph-based network analysis of RSFC data, particularly techniques derived from graph theory, has become widely used for analyzing brain network connectivity (Islam et al., 2018; Rubinov & Sporns, 2010; Sporns, 2015; Sporns, 2018). With this method, many studies have found alterations in neurological disorders (Aracil-Bolaños et al., 2022; Wolf et al., 2011; Xu et al., 2016). With regard to narcissistic personality, as far as we know, only one study has reported abnormal topology within the DMN in young male patients using this method (Cao et al., 2022). In addition, a study on patients with ASPD showed increased functional connectivity, primarily in the DMN, which was also analyzed via this technique (Tang et al., 2016). However, these studies were limited to a small sample size (i.e., ranging from 19 to 64) and focused solely on the DMN. Therefore, a more balanced exploration of multiple personality traits through graph-based network analysis could provide deeper insights into the shared and distinct neural mechanisms of these traits.

To address these gaps in the literature, the main objective of the current investigation is to identify and characterize the functionality of macro-networks in both narcissistic and antisocial personality. We hypothesize that abnormalities within the DMN, SN, and FPN will be predictive of both traits. Specifically, we expected regional topological measures (such as the betweenness and eccentricity), as well as global topological measures (such as global efficiency, local efficiency, and average path length) within the DMN, SN, and FPN, to predict both narcissistic and antisocial personality traits.

For what concerns the SN, previous studies have found alterated local and global mtetrics in schizophrenia (Iglesias-Parro et al., 2023), major depression (Noman et al., 2024), dementia (Nigro et al., 2022), insomnia (Ma et al., 2018), Parkinson’s disease (Huang et al., 2019), and drug addiction (Mansoory et al., 2022). In the same vein, previous studies demonstrated alterations in the FPN in many psychiatric disorders such as ADHD (Wang et al., 2019), and, and Parkinson’s disease (Huang et al., 2019). The DMN has also been found to be alterated in many psychiatric disorders (Langerbeck et al., 2023). Based on these results, we aim to predict the extended role of the DMN, SN, FPN to narcissistic and antisocial personality traits.

However, we also expect some differences in local and global metrics in the two disorders. Narcissistic individuals spend a lot of time reflecting on the self to compensate for self-esteem with fantasies of power and importance (Di Pierro et al., 2016) whereas antisocial behaviors are less reflective and more oriented toward others to exploit and manipulate (De Wit-De Visser et al., 2023). Indeed, Antisocial individual are typically impulsive and do not reflect on themselves or the consequences of their actions (De Wit-De Visser et al., 2023; Korponay et al., 2017). This may be reflected in differences in how the DMN is related to these disorders. We predict that higher narcissistic traits will be associated with increased functionality, whereas antisocial traits will show a decreased DMN functionality.

To further understand the communications between the main regions of the macro-networks and other regions of the brain, seed-based connectivity will be considered too. The aim of seed-based analysis was to understand how key hubs within the triple network identified in graph theory analyses, may influence other brain regions. By combining these two analyses, we hope to enhance our understanding of local and distributed abnormal brain functionality in individuals with narcissistic and antisocial traits. We predict to find results that are coherent with the ones detected with the topological analysis.

## MATERIALS AND METHODS

### Participants

We utilized data from the MPI-Leipzig Mind Brain-Body dataset, which is accessible through the OpenNeuro database (Accession Number: ds000221) (Babayan et al., 2020). This dataset encompasses MRI and behavioral data gathered from 318 participants who participated in the project conducted by the Max Planck Institute (MPI) of Human Cognitive and Brain Sciences in Leipzig. The study was carried out under the authorization of the ethics committee at the University of Leipzig (Protocol ID: 154/13-ff) (Babayan et al., 2019).

For the purposes of this study, we rigorously selected data from individuals who participated in the LEMON and Neuroanatomy & Connectivity Protocols according to specific selection criteria. These criteria encompassed medical eligibility for magnetic resonance sessions, availability of structural T1-weighted images and functional MRI, and completion of the Personality Styles and Disorders Inventory (PSDI), with a focus on the narcissistic and antisocial scales. Participants aged 70 years or older were excluded to minimize potential confounding effects of age-related alterations in brain functional connectivity or topography (Li et al., 2021; Qin & Basak, 2020; Veréb et al., 2023)

The final sample comprised 183 healthy participants, including 89 females and 94 males. PSDI scores for narcissistic traits (*t*=0.89, *p*=.37) and antisocial traits (*t*=0.89, *p*=.39) did not differ significantly between genders. The participants’ages ranged from 22 to 68 years, with a mean age of 32.58 years (SD = 14.04 years). Age distribution was also balanced between genders (*t*=-0.82, *p*=.41).

### Personality Styles and Disorders Inventory (PSDI)

To assess personality traits we used the PSDI (German version: Persönlichkeits-Stil-und Störungs-Inventar, PSSI), a validated self-report inventory that measures various personality styles and can provide insights into potential personality disorders, particularly when extreme scores are observed. Developed and revised by Kuhl and Kazén (2009), the inventory consists of 140 items organized into 14 subscales. In our study, we focused specifically on the narcissistic (Mean score = 12.14, SD = ± 4.76) and antisocial (Mean score = 12.92, SD = ± 4.40) subscales. The score of the two sub-scale are correlate (*r* = .44, *p* < .01).

The PSDI shows a robust network of theoretically coherent relationships with a large number of clinical and non-clinical behavioral characteristics (e.g., suicidality, depression, psychosomatic symptoms, the Big Five personality traits, and the sixteen personality factors). This extensive network of associations supports the construct validity of the inventory. Furthermore, the PSDI employs objective scoring procedures and statistical analyses, and its consistency coefficients (Cronbach’s alpha) range between α = .73 and .85. Test-retest coefficients vary between *r* = .68 and .83, indicating good test-retest reliability. The construct validity of the inventory is considered acceptable for both clinical and non-clinical behaviors (Kuhl & Kazén, 2009).

### MRI Data Acquisition

The MPI-Leipzig Mind Brain-Body dataset comprises quantitative T1-weighted, functional, resting state, and diffusion-weighted images acquired at the Day Clinic for Cognitive Neurology of the University Clinic Leipzig and the Max Planck Institute for Human and Cognitive and Brain Sciences (MPI CBS) in Leipzig, Germany (Babayan et al., 2019). For the purpose of our research, we only considered the T1-weighted images. Magnetic Resonance Imaging (MRI) was performed on a 3T Siemens MAGNETOM Verio scanner (Siemens Healthcare GmbH, Erlangen, Germany) with a 32-channel head coil. The MP2RAGE sequence consisted of the following parameters: sagittal acquisition orientation, one 3D volume with 176 slices, TR = 5000 ms, TE = 2.92 ms, TI1 = 700 ms, TI2 = 2500 ms, FA1 = 4°, FA2 = 5°, pre-scan normalization, echo spacing = 6.9 ms, bandwidth = 240 Hz/pixel, FOV = 256 mm, voxel size = 1 mm isotropic, GRAPPA acceleration factor 3, slice order = interleaved, duration = 8 min 22 s.

### fMRI Data Pre-processing

Data pre-processing was performed using CONN (version 2022) (Whitfield-Gabrieli & Nieto-Castanon, 2012), SPM 12, and the MATLAB Toolbox (version 2021b). First, the CONN’s default pre-processing pipeline uses SMP12’s default parameters. This pipeline encompassed several stages: functional realignment and unwarping, translation and centering, conservative functional outlier detection, direct segmentation and normalization of functional data (1 mm resolution), translation and centering of structural data, segmentation and normalization of structural data (2.4 mm resolution), and lastly, spatial smoothing of functional and structural data using an 8 mm Gaussian kernel. Subsequently, the denoising phase was conducted. The aim of this phase is to pinpoint and eliminate confounding variables and artifacts from the estimated BOLD signal. These factors arise from three distinct sources: the BOLD signal originating from masks of white matter or cerebrospinal fluid, parameters and outliers defined during the pre-processing step, and an estimation of the subjects’ motion parameters.

In addition, functional data were denoised using a standard denoising pipeline, including the regression of potential confounding effects characterized by white matter timeseries, cerebrospinal fluid (CSF) timeseries motion parameters and their first order derivatives, outlier scans (below 79 factors), session effects and their first order derivatives, and linear trends within each functional run, followed by bandpass frequency filtering of the BOLD timeseries between 0.008 Hz and 0.09 Hz. CompCor noise components within white matter and CSF were estimated by computing the average BOLD signal as well as the largest principal components orthogonal to the BOLD average, motion parameters, and outlier scans within each subject’s eroded segmentation masks. From the number of noise terms included in this denoising strategy, the effective degrees of freedom of the BOLD signal after denoising were estimated to range from 181.1 to 207 (average 205.3) across all subjects.

At the first-level analysis, ROI-to-ROI connectivity (RRC) matrices were estimated characterizing the functional connectivity between each pair of regions among 164 HPC-ICA networks and Harvard-Oxford atlas ROIs (Whitfield-Gabrieli & Nieto-Castanon, 2012). Functional connectivity strength was represented by Fisher-transformed bivariate correlation coefficients from a general linear model (weighted-GLM), estimated separately for each pair of ROIs, characterizing the association between their BOLD signal time-series (see Figure 1).

**Figure 1.**
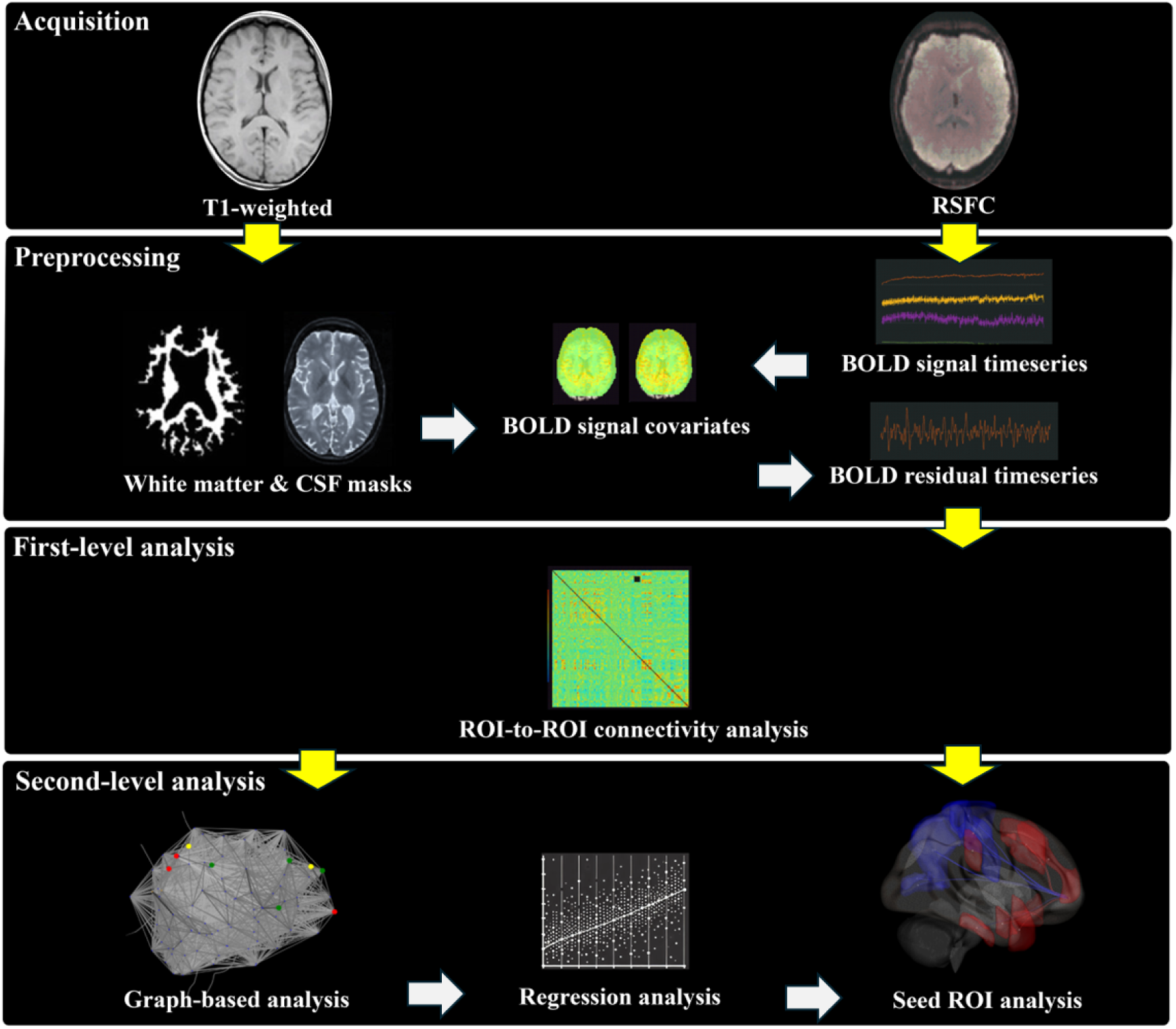
Data processing and analysis pipeline. Lines with an arrow indicate that the output of the previous step was supplied into the next step. The structural MRI (T1-weighted) and resting-state functional connectivity (RSFC) were acquired in the current study. The CONN toolbox was utilized to preprocess for both T1-weighted and RSFC. T1-weighted images were segmented into white matter and CSF masks. In the spatial domain, the BOLD signal time series was computed from the results of slice-timing, realignment, normalization, and smoothing. The BOLD residual timeseries were preprocessed at the temporal domain using the BOLD signal covariates, including segmented white matter and CSF masks, as well as BOLD signal timeseries. Analyses at the first-level were performed using the ROI-to-ROI connectivity approach. The results of this step were then used for subsequent analyses, including graph theory and seed ROIs.

### Graph Theory Analysis

The CONN toolbox was utilized to analyze graph theoretical metrics (Whitfield-Gabrieli & Nieto-Castanon, 2012). The toolbox characterizes structural properties of the estimated ROI-to-ROI functional connectivity networks and allows users to perform group-level analysis of these measures. Each subject-specific ROI-to-ROI connectivity matrix is thresholded at a fixed level (Nieto-Castanon, 2020; Whitfield-Gabrieli & Nieto-Castanon, 2012). Graph theory measurements depend on hypothesis levels, such as global measures, subnetwork levels measured by module, or region levels measured by node. In this study, we focused on regional measurements commonly used to assess nodal centrality in human brain functional connectivity, such as betweenness, closeness, eigenvector, and eccentricity (Boccaletti et al., 2006; Rubinov & Sporns, 2010; Zuo et al., 2011). Additionally, we examined global topology metrics, including global efficiency, local efficiency, average path length, and clustering coefficient, which have been previously identified in functional topological studies on NPD (Cao et al., 2022).

This approach differs significantly from classic random graph models in explaining the graph measurement results, particularly how nodes and edges are connected, influencing their contributions to network function. Network theory integrates the concept of functional specialization with that of distributed processing, thereby enhancing our understanding of brain function. By mapping brain connectivity, this approach reveals unique connectivity fingerprints for different regions, predicting their functional roles and highlighting the organization of functional groupings within the brain. Various centrality measures, such as degree and betweenness, are employed to identify influential network hubs and their relationships within the network’s community structure (Rubinov & Sporns, 2010; Sporns, 2018).

The prediction of narcissistic and antisocial trait scores was conducted using IBM SPSS version 25. In particular, multiple regression with a stepwise method was used, and predictors were included in global and regional metrics resulting from graph network analysis including the DMN the SN, and the FPN, as well as the visual and sensorimotor networks as control networks. Plotting a three-dimensional (3D) brain picture utilizing graph network results that identified ROIs or nodes that can predict narcissistic and antisocial behaviors was done using the CONN toolbox. Two-sided correlations and uncorrected p <.05 were used to threshold the 3D-rendered brain view of the analyzed network of functional connectivity.

Seed ROI analysis was performed using the CONN toolbox. We examined the between-subject effects of narcissistic and antisocial scores to establish connections between whole ROIs and constructed network matrices that resulted from the previous step. The individual ROI maps were generated to include threshold ROI-to-ROI connections based on intensity, applying a two-sided threshold for negative and positive seed levels. The significance of seed ROIs was determined through uncorrected *p* < .05. Finally, the survival ROIs of each seed region were visualized in 3D brain plots using the CONN toolbox. The 3D-rendered view figures show the supra-threshold ROI-level findings that the connectivity contrast effect sizes (between the seed and each target) and uncorrected *p*-values for the designated second-level analysis were shown for each target ROI. These connectivity directions and intensity were demonstrated by the negative color in blue and the positive color in red (Nieto-Castanon, 2020).

## RESULTS

### Graph-based network results

#### Narcissistic traits

The regression results indicated that the regional topological model accounted for a significant proportion of variance in narcissistic traits, *F*(4,95)=6.96, *p*<.001 F(4,95)=6.96, *p*<.001. Specifically, the eigenvector centrality of the MPFC within the DMN was a significant positive predictor (*β* = 0.17, *t*=2.29, *p*=.02), while the eccentricity of the ACC region in the SN was a significant negative predictor (*β*=−0.17, *t*=−2.35, *p*=.02). Additionally, the betweenness centrality of the MPFC in the DMN (*β*=0.15, *t*=2.06, *p*=.04) and the betweenness centrality of the left anterior insula in the SN (*β*=−0.15, *t*=−2.06, *p*=.04) were also significant predictors (See Table 1).

**Table 1.**
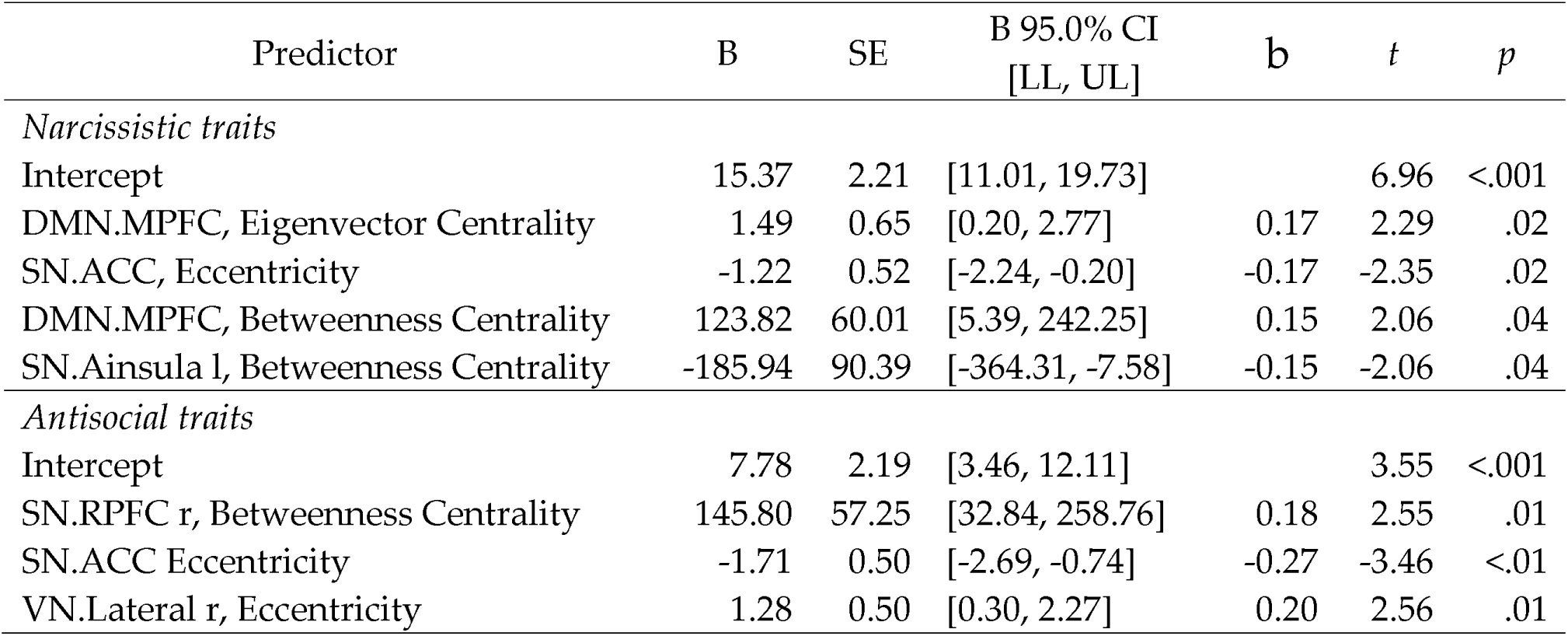
Multiple regression analysis in regional topological matrices predicting narcissistic and antisocial traits.

Furthermore, the overall global topological model was found to be significant, *F*(2,97)=4.87, *p*<.01. Although the intercept was not significant (*t*=−1.272, *p*=.21), the global efficiency of the MPFC in the DMN significantly predicted narcissistic traits (*β*=0.16, *t*=2.15, *p*=.03), as did the local efficiency of the left LPFC in the FPN (*β*=0.15, *t*=2.08, *p*=.04) (See Table 2).

**Table 2.**
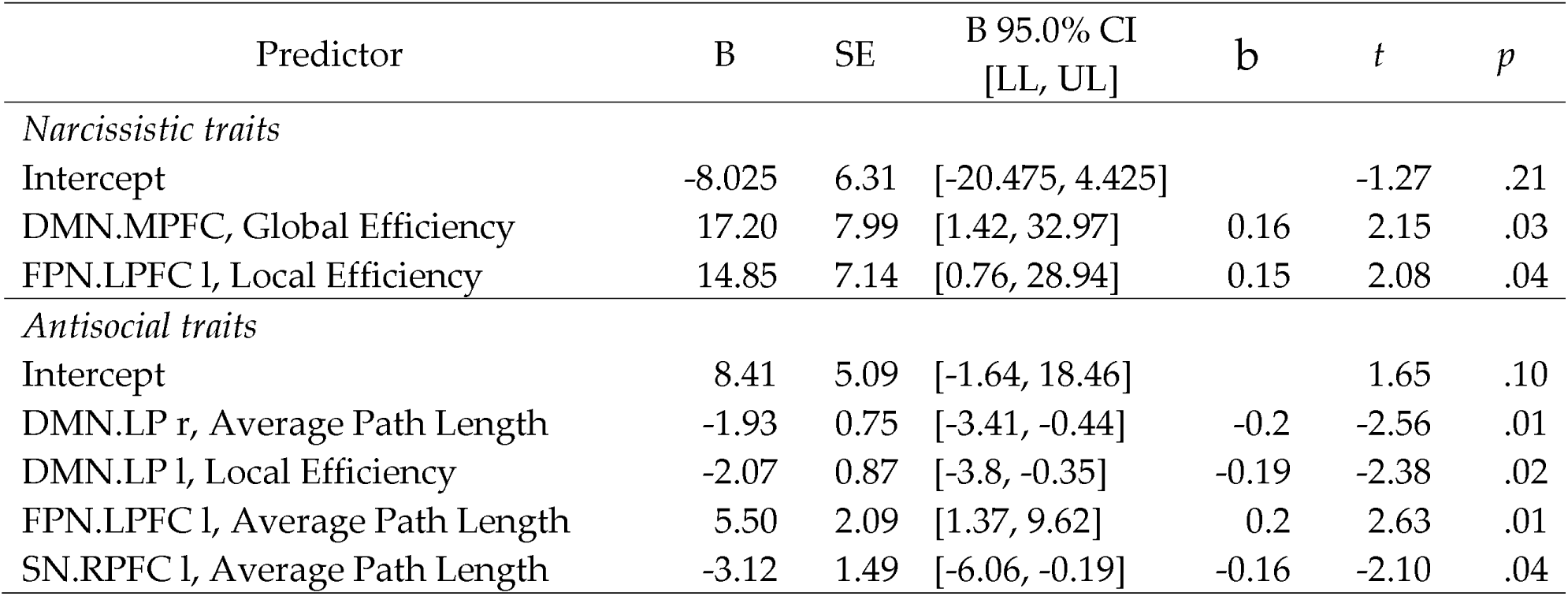
Multiple regression analysis in global topological matrices predicting narcissistic and antisocial traits.

#### Antisocial traits

The results of the regression indicated that the regional topological model explained a significant proportion of the variance in antisocial traits, F(3,96)=4.92,p<.001. The betweenness centrality of the right RPFC within the SN was found to be a significant positive predictor (*β*=0.18, *t*=2.55, *p*=.01), while the eccentricity of the ACC in the SN was found to be a significant negative predictor (*β*=−0.27, *t*=−3.46, *p*<.01). Additionally, the eccentricity of the right lateral part in the VN was also a significant positive predictor (*β*=0.20, *t*=2.56, *p*=.01), see Table 1.

Likewise, the global topological model was significant, *F*(4,95) = 5.73, *p*<.001. Although the intercept was not significant (*t*=1.65, *p*=.10), the average path length of the right LP in the DMN (*β*=−0.20, *t*=−2.56, *p*=.01) and the local efficiency of the left LP in the DMN (*β*=−0.19, *t*=−2.38, *p*=.02) were significant negative predictors. Conversely, the average path length of the left LPFC in the FPN (*β*=0.20, *t*=2.63, *p*=.01) and the average path length of the left RPFC in the SN (*β*=−0.16, *t*=−2.10, *p*=.04) were significant predictors, see Table 2.

**Figure 2.**
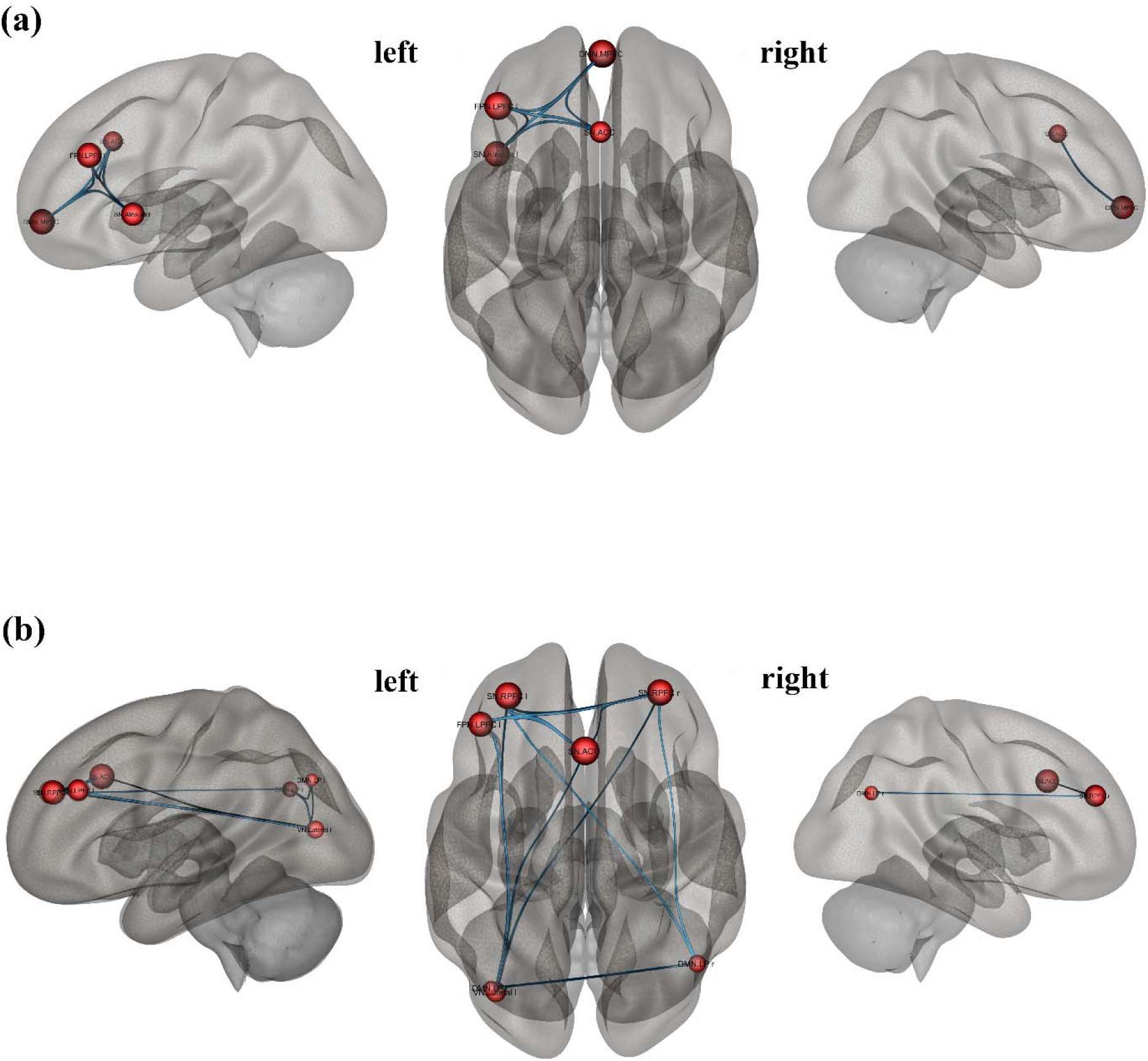
presents the brain network from graph-based analysis, providing a 3D representation of the brain’s connectivity concerning narcissistic traits (a) and antisocial traits (b). Seed ROIs are presented as nodes in red while two-side correlations between nodes are presented in blue lines (*p*_unc_ <.05).

### Seed ROI results

When analyzing network metrics to predict narcissistic and antisocial traits, four key nodes were identified as significant predictors: DMN.MPFC, SN.ACC, SN.Ainsula l, and FPN.LPFC l. The following sections present the seed ROI results, which demonstrate that ROI targets were related to these seed sources/nodes, with a significant between-subject effect on narcissistic and antisocial scores.

#### Narcissistic traits

The seed regions and targets included ROIs that showed significant connections for narcissistic scores at an uncorrected p-value of 0.05, see Table 2. Firstly, the MPFC of the DMN showed significant positive associations with multiple brain regions, including the left temporal pole, right frontal orbital cortex, and posterior inferior temporal gyrus on the left side (all p_unc_ < .01). Other positively associated areas (p_unc_ ranging from .02 to .04), included regions such as the right and left lateral prefrontal cortex, left supramarginal gyrus, and right middle frontal gyrus. Conversely, significant negative associations were found with regions such as the superior SMN, right supracalcarine cortex, and precuneus cortex (all p_unc_ = .01 or .02), as visualized in Figure 3 (a). Secondly, the ACC in the SN revealed significant positive associations with regions such as the right paracingulate gyrus and the right temporooccipital middle temporal gyrus (p-unc < .01). Other regions, such as the posterior superior temporal gyrus on the left side and the right frontal pole, also showed positive associations (p_unc_ around .02). Additionally, the left superior frontal gyrus exhibited a significant positive association (p_unc_ = .04), as shown in Figure 4 (a). Thirdly, the Ainsula in the SN demonstrated significant positive associations with the left frontal lobe and the DMN medial prefrontal cortex (p_unc_ = 0.01 and 0.02, respectively), highlighting its specific connectivity and interaction with the DMN and other prefrontal regions, as visualized in Figure 5. Lastly, the LPFC in the FPN showed significant positive associations with several regions, including the right anterior superior temporal gyrus and the left anterior temporal fusiform cortex (p_unc_ < .01). Additional positive associations were found with areas such as the DMN medial prefrontal cortex, left temporal pole, and right Heschl’s gyrus (p_unc_ ranging from .02 to .04). On the negative side, significant associations were observed with regions like the left juxtapositional lobule cortex and left cerebellum Crus2 (p_unc_ <.01), as well as other areas like the superior SMN and Vermis 7 (p_unc_ = 0.02 to 0.04), as visualized in Figure 6 (a).

**Figure 3.**
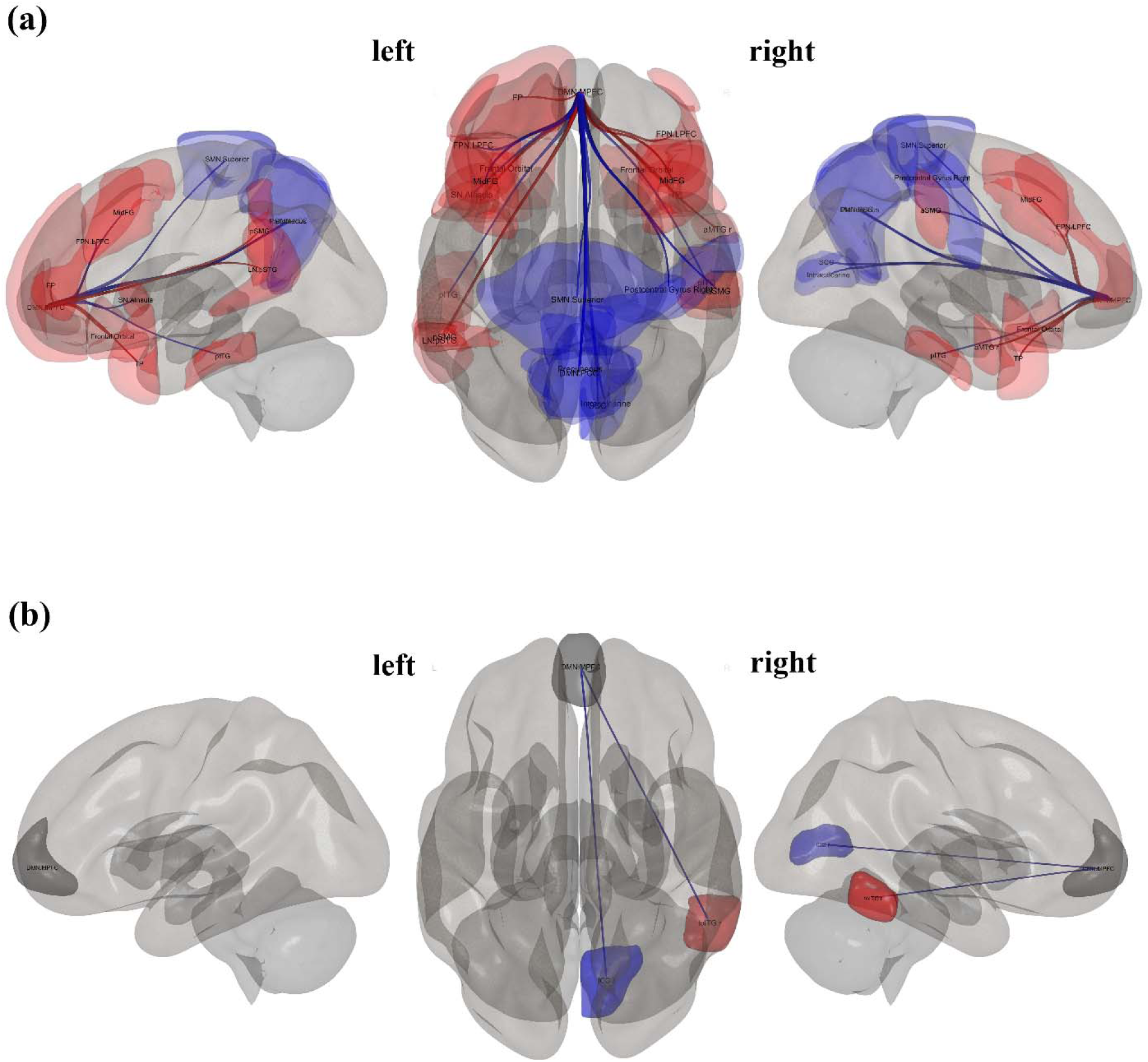
presents 3D brain plots, specifically the seed ROI from DMN.MPFC, which is known to correlate with other regions (*p*_unc_ <.05). The positive correlation in red lines and negative correlation in blue lines indicate the strength and direction of these correlations, providing a visual representation of the brain’s activity in relation to narcissistic traits (a) and antisocial traits (b).

**Figure 4.**
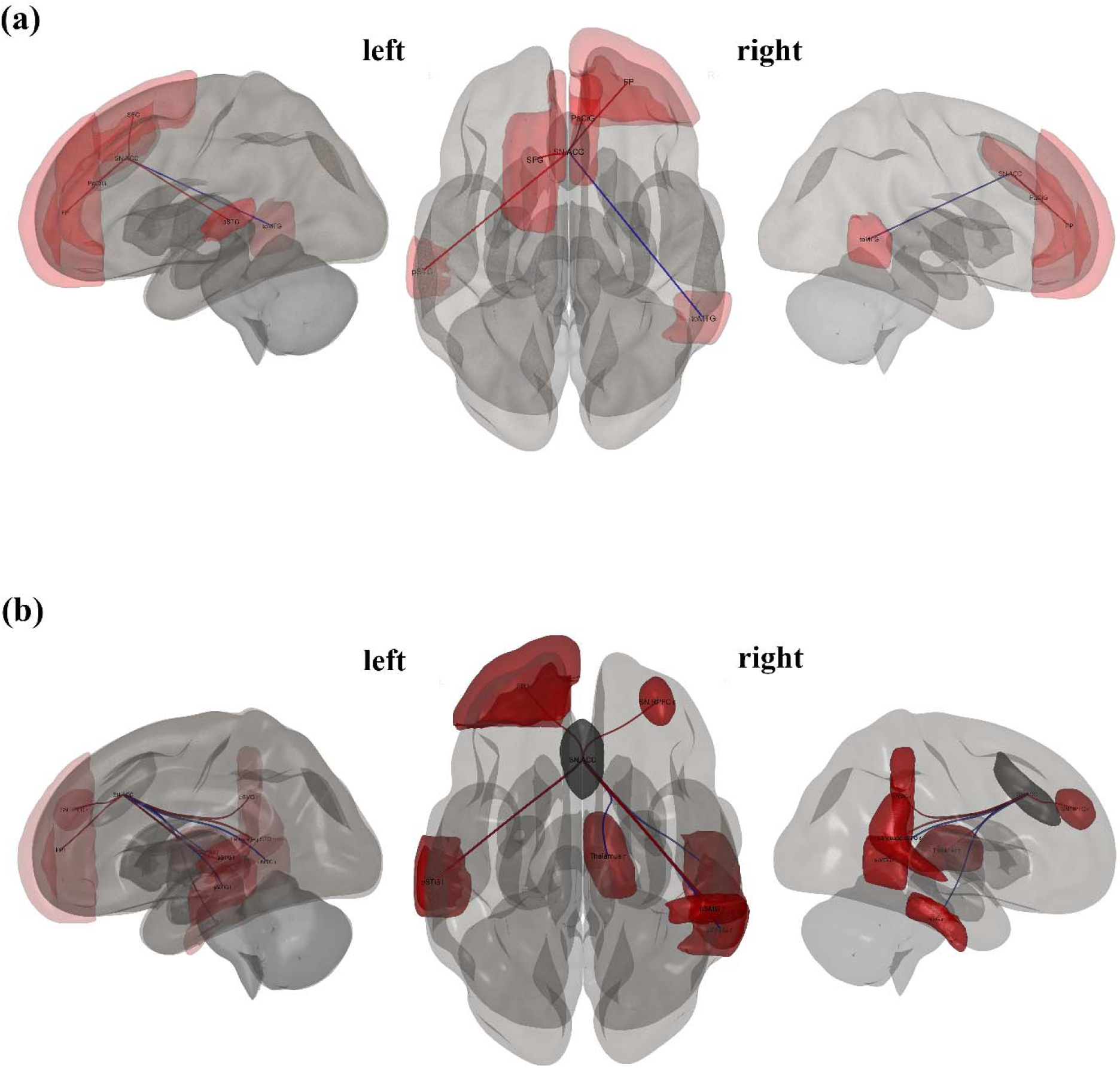
presents 3D brain plots, specifically the seed ROI from SN.ACC, which is known to correlate with other regions (*p*_unc_ <.05). The positive correlation in red lines and negative correlation in blue lines indicate the strength and direction of these correlations, providing a visual representation of the brain’s activity in relation to narcissistic traits (a) and antisocial traits (b).

**Figure 5.**
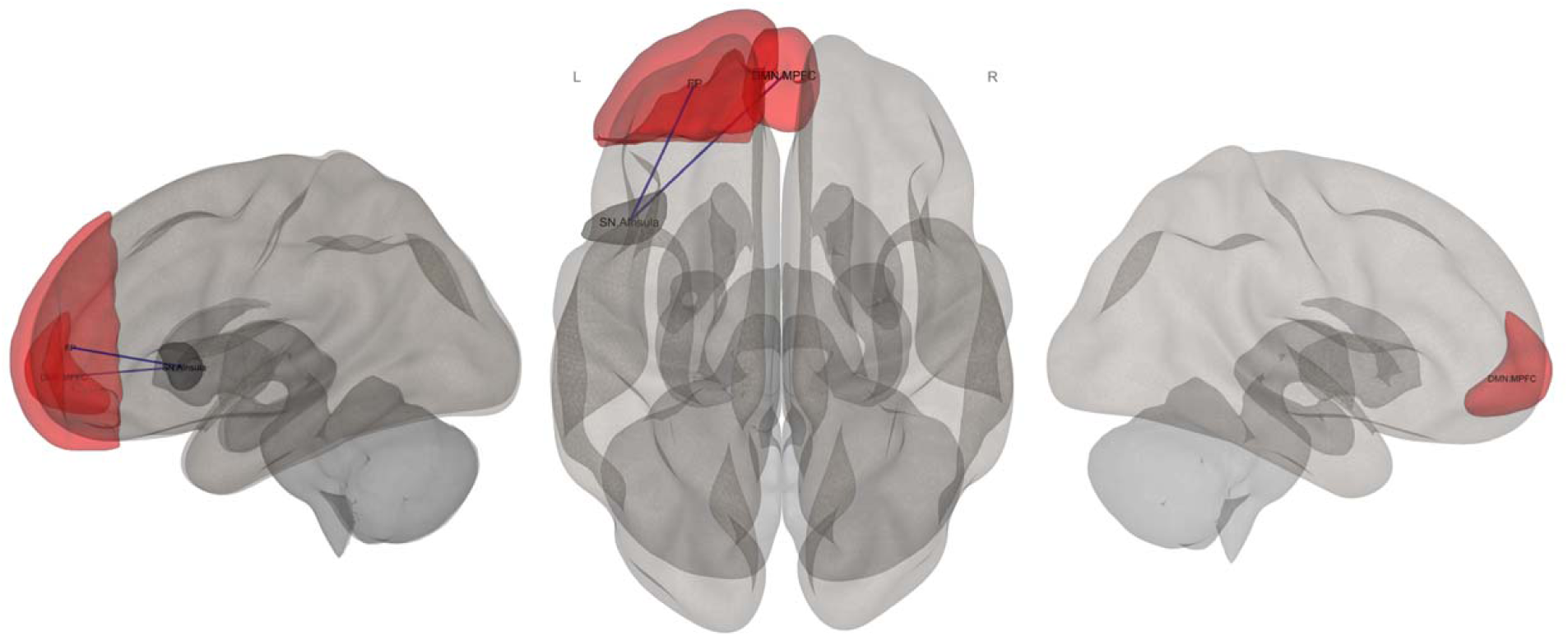
presents 3D brain plots, specifically the seed ROI from SN.AInsula left, which is known to correlate with other regions (*p*_unc_ <.05). The positive correlation in red lines and negative correlation in blue lines indicate the strength and direction of these correlations, providing a visual representation of the brain’s activity in relation to narcissistic traits.

**Figure 6.**
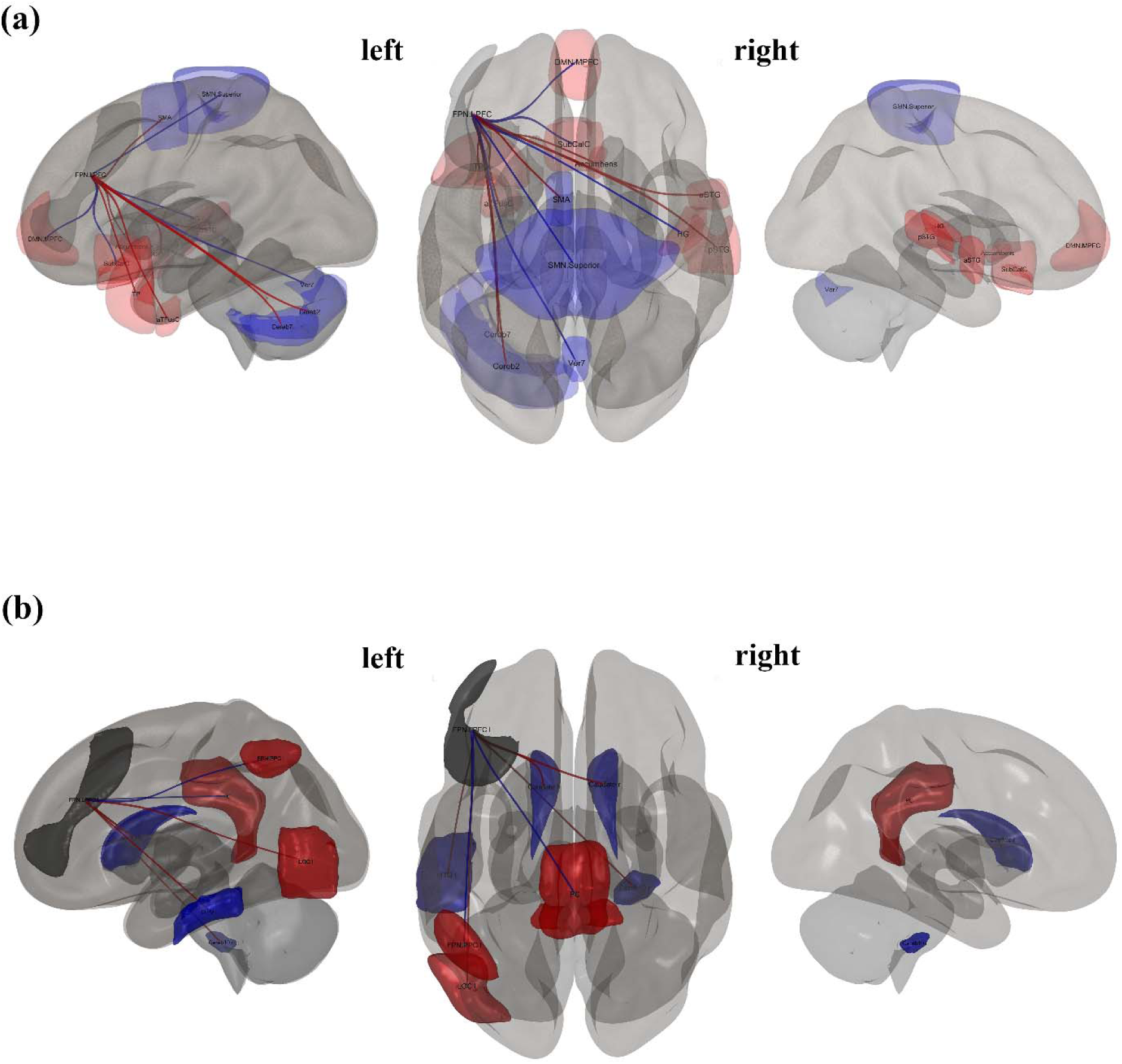
presents 3D brain plots, specifically the seed ROI from FPN.LPFC, which is known to correlate with other regions (*p*_unc_ <.05). The positive correlation in red lines and negative correlation in blue lines indicate the strength and direction of these correlations, providing a visual representation of the brain’s activity concerning narcissistic traits (a) and antisocial traits (b).

#### Antisocial traits

The seed regions and targets included ROIs that show significant connections for antisocial scores at an uncorrected p-value of 0.05, see Table 4. The MPFC of the DMN showed significant positive associations with the Inferior Temporal Gyrus, temporooccipital right (p-unc = .01) and significant negative associations with the Intracalcarine Cortex, right (p_unc_ = .03), as visualized in Figure 3 (b). Similarly, the ACC in the SN demonstrated significant positive associations with the LN Superior Temporal Gyrus, posterior right (p-unc = .01) and the SN Rostral Prefrontal Cortex, right (p_unc_ = .02), among others, with p_unc_ values ranging from .02 to .04 for additional targets, as visualized in Figure 4 (b). Conversely, the left LPFC in the FPN revealed significant negative associations with the Cerebellum 10, Right (p_unc_ = .01) and positive associations with the Lateral Occipital Cortex, inferior left (p_unc_ = .02), while also encompassing targets with p_unc_ values of .04, as visualized in Figure 6 (b). The left RPFC in the SN exhibited significant negative associations with the Cingulate Gyrus, posterior (p_unc_ < .01) and positive associations with the right Amygdala (p_unc_ = .02), while also including targets with p_unc_ values of .04. On the other hand, the right RPFC in the SN showed significant positive associations with the anterior left Temporal Fusiform, (p_unc_ < .01) and several other regions with p_unc_ values ranging from .02 to .04, as visualized in Figure 7 (a). Lastly, the left LP in the DMN showed significant negative associations with the left Cerebellum 9 (p_unc_ < .01) and a positive association with the SN.RPFC (p_unc_ = .04). In opposition, the right LP in the DMN exhibited significant positive associations with the right SN.SMG (p_unc_ = .01), the right Parietal Operculum (p_unc_ = .01), and the left Lingual Gyrus (p_unc_ = .01). Additionally, regions such as the LN.pSTG and toMTG r showed p_unc_ values of .03, while the FPN.LPFC and the pSTG r (p_unc_ = .04) were also associated (p_unc_ = .04), as visualized in Figure 7 (b).

**Table 3.**
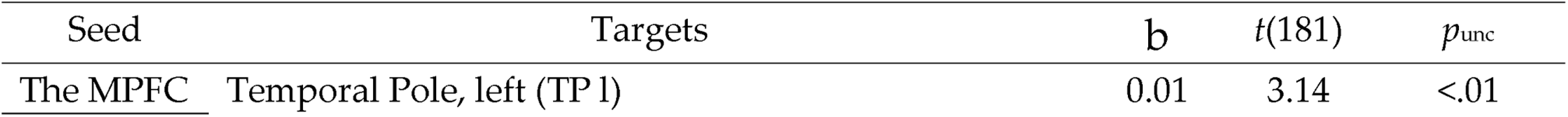
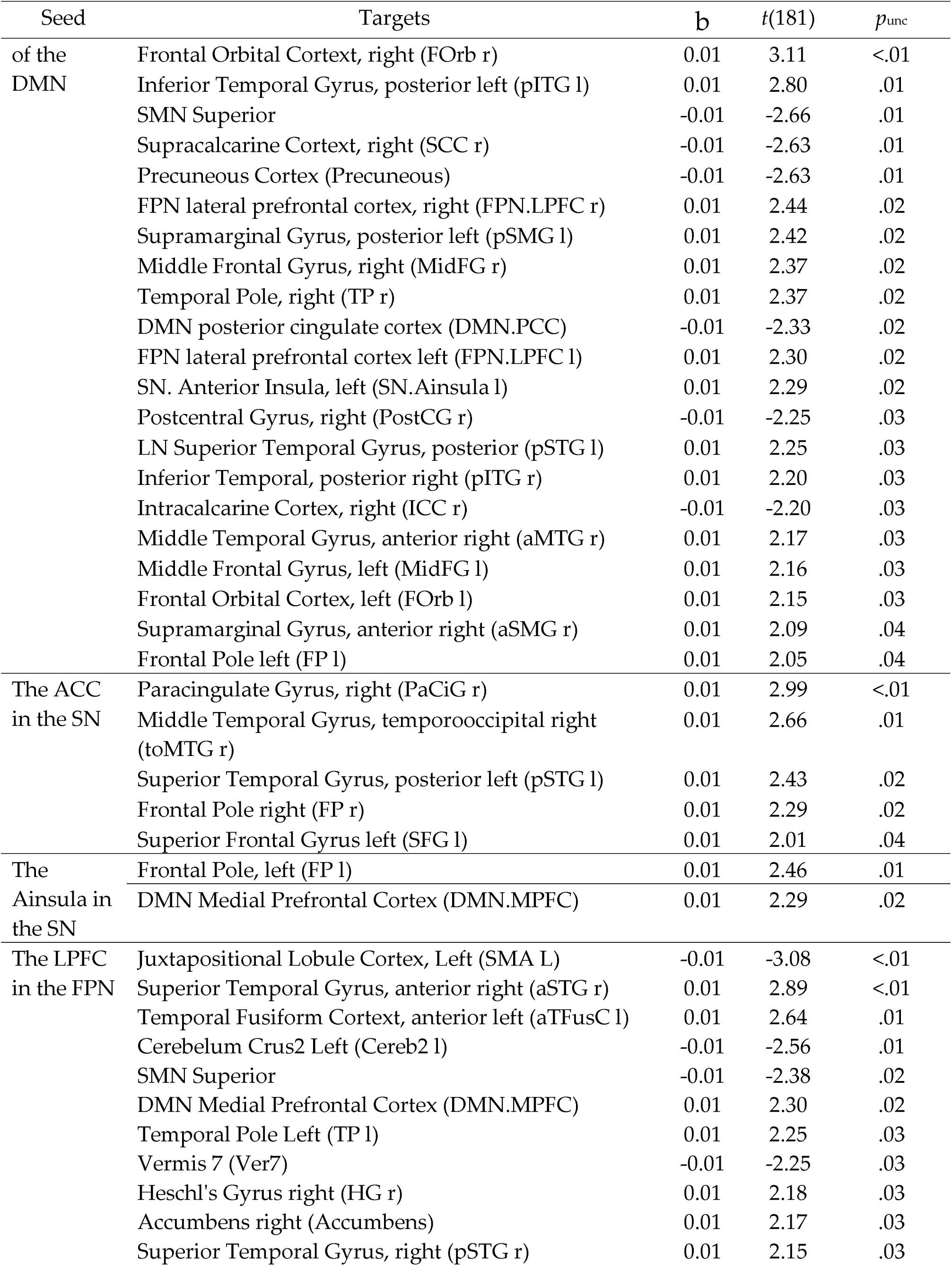
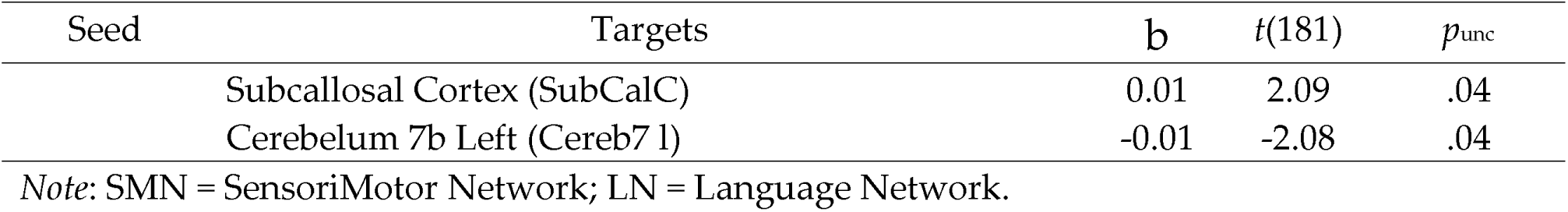
The seed regions and targets included ROIs that were significantly associated with narcissistic scores at an uncorrected p-value of 0.05.

**Figure 7.**
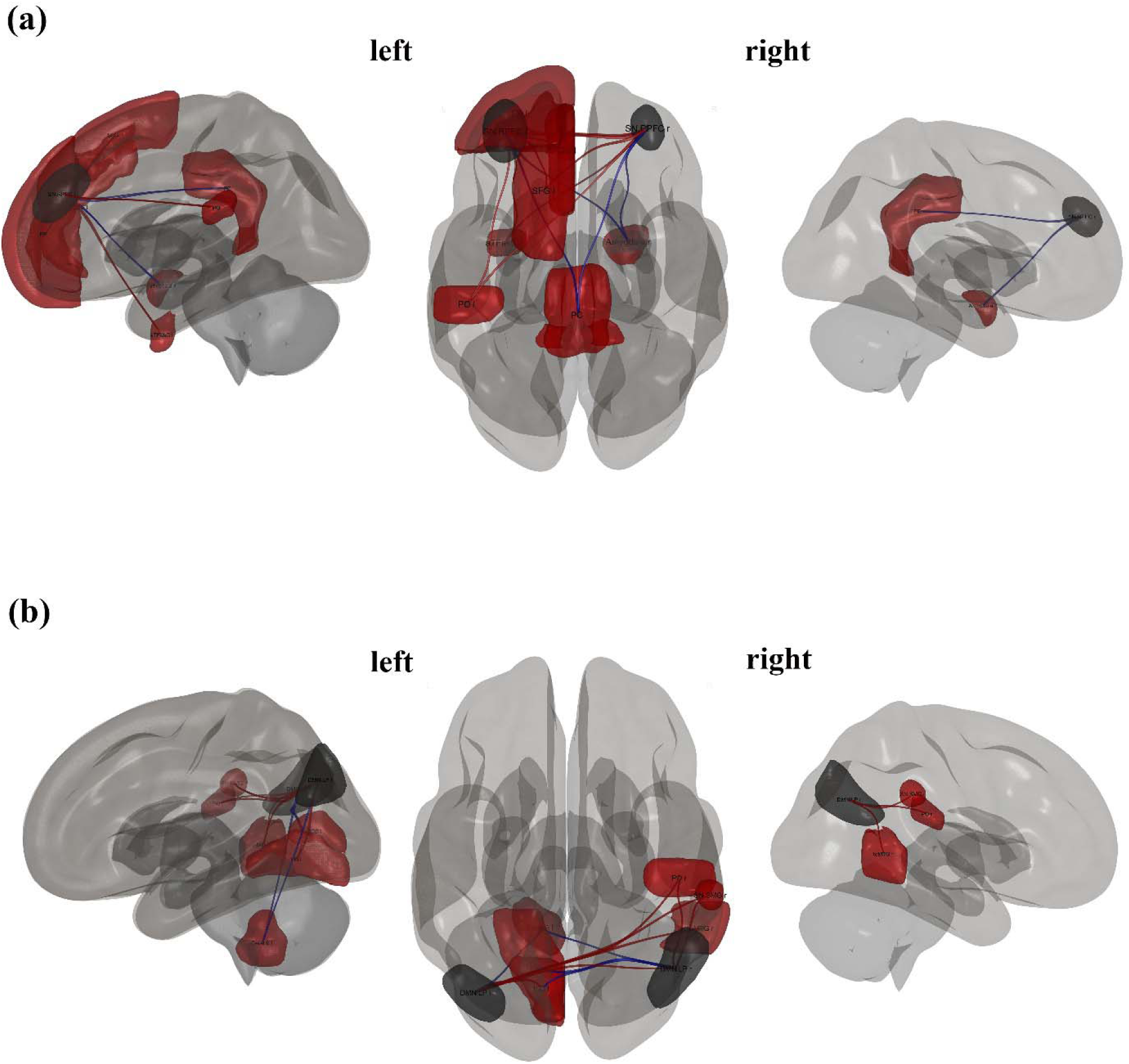
presents 3D brain plots, specifically the seed ROI from SN.RPFC (a) and DMN.LP (b), which is known to correlate with other regions (*p*_unc_ <.05). The positive correlations are represented in red lines, and the negative correlations in blue lines, indicating the strength and direction of these associations, providing a visual representation of the brain’s activity in relation to antisocial traits.

**Table 4.**
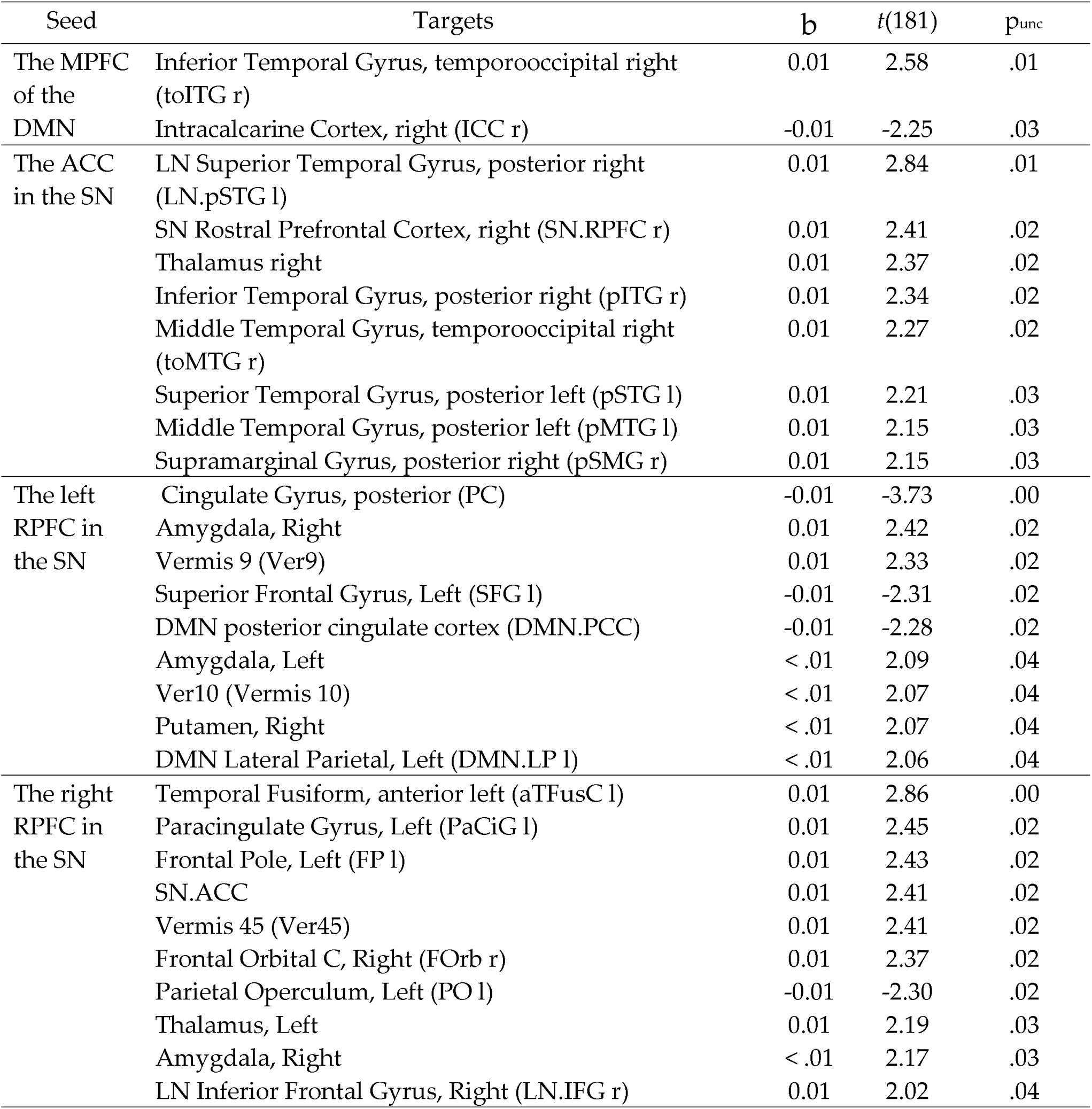
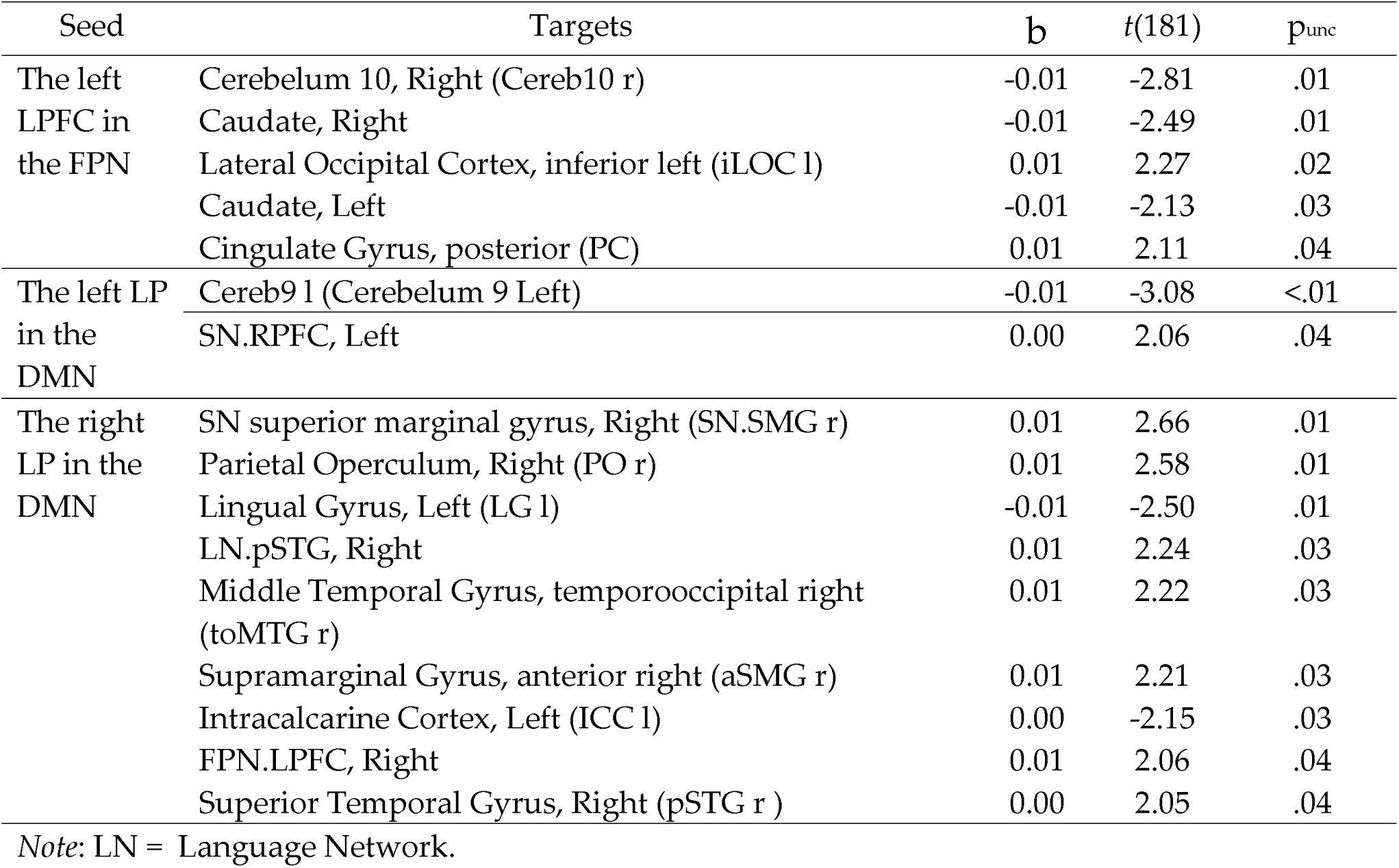
The seed regions and targets included ROIs that were significantly connected for antisocial scores at a 0.05 uncorrected p-value.

## DISCUSSION

In our study, we employed a graph-based network approach to identify macro-network functionalities that could elucidate the complex connectivity patterns associated with narcissistic and antisocial traits. We hypothesized that global and regional measures of the DMN, SN, and FPN (i.e., the “triple network”) could predict both traits. This could provide evidence that these two personality types share not only symptoms but also similar brain characteristics.

As predicted, both traits were associated with reduced intra-network connectivity within the SN, particularly in the ACC, and increased local efficiency (more effective processing) in the LPFC of the FPN, indicating similar altered patterns of emotional and cognitive control. Although we also confirmed a role for the DMN, this network displayed notable differences between the two traits. Narcissistic personality traits (NPT) were linked with increased DMN activity, especially in the medial prefrontal cortex (MPFC), while antisocial personality traits (APT) showed decreased DMN involvement. Higher betweenness centrality in the right RPFC of the SN and increased VN activity were observed only for APT.

These findings highlight both shared and distinct neural underpinnings, with the DMN’s self-reflective processes being more related to narcissism and the SN’s emotional processes being more critical for antisocial traits. In the following sections, we will discuss these results in detail.

### Shared Mechanisms

At a regional topological level, we found that the eccentricity measure of the ACC within the SN was predictive of both traits. Eccentricity refers to how much a node (in this case, the ACC) is connected to other nodes within the network (SN). The fact that eccentricity negatively predicts both traits indicates that the higher the personality traits, the lower the connectivity between the ACC and other regions of the SN. In other words, abnormal intra-network connectivity in the SN is associated with both traits. This result aligns with previous evidence showing that altered GM in the ACC predicts narcissistic traits (Jornkokgoud et al., 2024; Jornkokgoud et al., 2023). We also found that other hubs of the SN were affected in both traits, but with some differences. Specifically, the anterior insula (AI) of the SN was predictive of narcissistic traits, whereas the RPFC hub was more predictive of antisocial traits. Overall, the AI and the ACC are considered central nodes within the SN, known to respond to behaviorally salient events by integrating them with the emotional context (Jankowiak-Siuda & Zajkowski, 2013; Li et al., 2018; Sridharan et al., 2008). The AI and ACC are also active in response to experiences related to empathy, social rejection, high anxiety, or low self-esteem, playing a major role in self- and other-related emotional processing—especially relevant in the context of narcissistic traits (Cascio et al., 2014; Jankowiak-Siuda & Zajkowski, 2013; Jauk & Kanske, 2021). Although the AI was not identified as a predictor of antisocial traits in our results, previous research has found reductions in GM volume and increased effective connectivity in the AI, as well as abnormal activation in individuals with antisocial behavior and a thinner-than-normal cortex in those with psychopathy (Aoki et al., 2014; Sitaram et al., 2014). In addition, the AI was highlighted as significantly involved in individuals with ASPD with psychopathy compared to those without psychopathy and non-offenders, indicating distinct neural activation patterns between subtypes of ASPD (Gregory et al., 2015).

Regarding global topological metrics, we found that local efficiency in the lateral prefrontal cortex (LPFC) of the FPN was associated with both disorders. Higher efficiency in this region within the FPN correlated with higher narcissistic and antisocial traits. The LPFC within the FPN is essential for executive control, integrating cognitive processes necessary for purposeful behavior. This region regulates attentional control, filters relevant information from distractions, and facilitates action selection to maintain focused attention and select behavioral targets (Tanji & Hoshi, 2008). Additionally, the LPFC and MPFC are involved in higher-order cognitive functions such as decision-making and action planning, which are critical for the adaptive control of behavior (Nee & D’Esposito, 2017; Tanji & Hoshi, 2008). This result may indicate a greater capacity for planning and decision-making, aiding in the pursuit of self-serving goals, which is characteristic of both narcissistic and antisocial individuals.

We also found DMN involvement for both traits, although with opposite patterns ( increased activity for narcissism and decreased activity for antisociality), and different subregions implicated (MPFC for narcissism and LPFC for antisociality). This suggests that both traits are encoded within the DMN, but with notable differences. The overexpression of the DMN in individuals with high narcissistic traits aligns with prior findings in other personality disorders (Grecucci et al., 2022; Grecucci et al., 2023; Langerbeck et al., 2023) and may underlie tendencies for overthinking and distorted self-representations (Steiner et al., 2021; Wang et al., under review). In contrast, reduced DMN activity in individuals with antisocial traits may reflect a diminished capacity for self-reflection and more externally oriented thinking (Kılıçaslan et al., 2022).

Regarding the seed ROI analysis, the results showed that both narcissistic and antisocial traits involve overlapping regions within the SN and FPN, with significant connections, but not within the DMN. In the MPFC of the DMN, we found that connectivity associated with narcissistic traits significantly correlated with areas including the frontal, temporal, and cingulate cortices, while antisocial traits were primarily connected to regions in the temporal and occipital areas. These findings are consistent with our topological findings. Furthermore, the ACC within the SN was found to predict both traits, connecting with regions such as the right PaCiG, STG, right MTG, and right frontal lobe in relation to narcissism, while in antisocial traits, it connected with similar regions, including the temporal gyrus and the right thalamus. These results are consistent with previous findings showing the ACC’s involvement in emotional regulation and reward processing (Dugré & Potvin, 2022; Flannery et al., 2020). In contrast, the LPFC of the FPN was implicated in both traits, with correlations found between this region and the prefrontal cortex, motor cortex, and cerebellum. In narcissism, the LPFC was connected with the STG and anterior fusiform cortex (aTFusC), while in antisocial traits, was connected with the cerebellum and cingulate gyrus. Previous evidence suggests that LPFC connectivity is crucial for cognitive control (Nee & D’Esposito, 2017). Specifically, individuals with antisocial traits exhibit sub-optimal FPN topology, likely linked to inefficient neural communication, which may contribute to difficulties in executive functioning (Lumaca et al., 2024). Our findings are consistent with prior studies demonstrating that GM and WM in regions like the MidFG, STG, and cerebellum contribute to the prediction of narcissistic traits (Jornkokgoud et al., 2024; Jornkokgoud et al., 2023).

### Distinct Mechanisms

The DMN (particularly the MPFC hub) was found to be more involved in narcissistic traits, but not in antisocial traits. This finding aligns with the differences in self-reflective behavior between these two traits. Previous studies have suggested that narcissistic traits are reflected in the GM and WM networks contributing to the DMN (Jornkokgoud et al., 2024; Jornkokgoud et al., 2023). Specifically, earlier research indicated that cortical volume in the MPFC within the DMN is negatively correlated with pathological narcissism (Mao et al., 2016). The MPFC is crucial for social cognition, which includes recognizing and understanding mental states, intentions, and emotions—both in oneself and in others (Le Petit et al., 2022). Abnormalities in cortical volume in this region are linked to deficits in these cognitive functions, which are essential for empathy and theory of mind (Mao et al., 2016; Massey et al., 2017).

In contrast, higher betweenness centrality in the right RPFC of the SN was solely likely to antisocial traits. RPFC is involved in higher-level cognition, such as multitasking, goal maintenance, and integrating information from internal and external sources. It helps balance focus between external stimuli and internal thoughts, such as future planning (Benoit et al., 2011). This may explain why individuals with antisocial traits, who generally display poor impulse control and a tendency to act without thinking (Kernberg, 1992; Miller et al., 2017), have lower self-regulation compared to narcissistic individuals.

Another notable difference between the two traits was the presence of the visual network in antisocial individuals, bit not in narcissistic individuals. Recent studies, such as Bakiaj et al. (2024), have suggested that antisocial individuals may display heightened visual abilities. Similarly, Espinoza et al. (2018) demonstrated that increased functional connectivity in the visual network, along with the SN and DMN, is associated with individuals displaying antisocial behavior, particularly forensic populations (Espinoza et al., 2018). However, earlier studies like Philippi et al. (2015) did not find associations between the visual and auditory networks and psychopathy in prison inmates (Philippi et al., 2015). As expected, the sensorimotor network, which was used as a control in this study, did not predict either narcissistic or antisocial traits.

Our seed ROI results further showed that both narcissistic and antisocial traits involve different regions within the DMN and SN. In individuals with narcissistic traits, the MPFC region of the DMN was associated with various targets, including the temporal pole, frontal orbital cortex, inferior temporal gyrus, and precuneus. These findings are consistent with earlier research showing that abnormalities in the GM volume of the MPFC are linked to reward and addiction, as well as socio-emotional processes such as empathy and emotional regulation (Myznikov et al., 2024; Pastor & Medina, 2021; Rolls et al., 2020). Although the MPFC is also involved in antisocial traits, its connectivity pattern differs. In antisocial individuals, it connects with regions such as the right inferior temporal gyrus and intracalcarine cortex, which are associated with visual and sensory processing. This may indicate altered responses to social cues. Additionally, the anterior insula and ACC within the SN are more related to narcissistic traits, with these regions showing connections to areas like the paracingulate gyrus and superior temporal gyrus. These results align with the topological metrics discussed earlier and are supported by recent studies suggesting reduced amygdala functioning as a central hub in the global information flow in psychopathic individuals (Tillem et al., 2019).

### Translational Implications

Our findings suggest common and distinct neural correlates for narcissism and antisociality, offering important insights for clinical practice, particularly in diagnostics and treatment planning. For example, DMN hubs were negatively correlated with antisocial traits and positively correlated with narcissism, possibly explaining the differences in self-reflection and sense of self between the two traits. This could lead to improved diagnostics for NPD and ASPD, which are often comorbid (Kraus & Reynolds, 2001; Widiger, 2011). The similarities in brain connectivity in the SN and FPN for both traits may be linked to the fact that individuals with these traits tend to exhibit dangerous behaviors and decreased awareness of risks, along with improved strategic planning and manipulation to achieve their goals. Moreover, the moderate relationship between narcissistic and antisocial subscales indicates overlapping characteristics in terms of the PSDI. These suggest that the dimensions and items of narcissistic and antisocial personality tests should be correlated and could be used to design more targeted interventions for individuals with these personality disorders, tailored to their specific neural profiles.

### Conclusions and limitations

In the present study, we found a clear involvement of the DMN, SN, and FPN in narcissistic and antisocial traits expanding previous evidence of their role in other personality disorders (Cao et al., 2022; Feng et al., 2018; Yang et al., 2015). The DMN’s involvement in self-referential and introspective processes, the SN’s role in detecting and integrating salient stimuli, and the FPN’s role in regulating self-oriented control underscore the complex interplay of these networks in shaping narcissistic and antisocial personality traits.

Our research does not come without limitations. First, this study only focused on resting-state functional MRI data, and future research could explore the fusion of functional and structural MRI data, such as gray or white matter. Second, The PSSI was used in our study to evaluate narcissism, and it did not distinguish between the grandiose and vulnerable subtypes. Future research on this issue should go deeper and explore potential distinctions between the two forms of narcissism in terms of the triple brain network. Third, although we used the largest sample to date to our knowledge, future studies may benefit from increasing the sample size to obtain more robust results. Lastly as gender differences were not the focus of this study, we excluded gender effects from our analyses to isolate the influence of personality traits on connectivity measures. Future studies could investigate potential gender differences in these personality traits.

In conclusion, our study confirms the involvement of the triple network in narcissistic and antisocial personalities, highlighting both similarities and differences, thereby advancing our understanding of this topic. These findings could inform the development of clinical interventions aimed at ameliorating these traits by targeting the functionality of the networks found to be altered.

## DATA AVAILBILITY STATEMENT

The data that support the findings of this study are openly available in OpenNeuro at https://doi.org/10.18112/OPENNEURO.DS000221.V2, reference number ds000221.

